# A trade-off in controlling upstream and downstream noise in signaling networks

**DOI:** 10.1101/2023.08.29.555248

**Authors:** Ka Kit Kong, Chunxiong Luo, Feng Liu

## Abstract

Signal transduction, underpinning the function of a variety of biological systems, is inevitably affected by fluctuations. It remains intriguing how the timescale of a signaling network relates to its capability of noise control, specifically, whether long timescale can average out fluctuation or accumulate fluctuation. Here, we consider two noise components of the signaling system: the upstream noise from the fluctuation of the input signal and the downstream noise from the stochastic fluctuations of the network. We discover a fundamental trade-off in controlling the upstream and downstream noise: a longer timescale of the signaling network can buffer upstream noise, while accumulate downstream noise. Moreover, we confirm that this trade-off relation exists in real biological signaling networks such as a fold-change detection circuit and the p53 activation signaling system.

**Author Summary:** Information transmission is vital in biological systems, such as decoding the information regarding nutrient levels during chemotaxis or morphogen concentrations in tissue development. While fluctuations arising from the stochastic nature of biological processes inevitably affect information transmission, noise control mechanisms have been studied for decades. However, it remains controversial what the role of the slow dynamics (long timescale) is in noise control. On the one hand, it has been reported to attenuate noise by averaging out fluctuations. On the other hand, it is also proposed to amplify noise by accumulating fluctuations. Here we dissect the noise in signaling systems into two components: upstream noise originating from signal fluctuation, and downstream noise from the network stochasticity. Our analysis reveals that upstream noise negatively correlates with timescale, while downstream noise exhibits a positive correlation, indicating a fundamental trade-off in controlling the two noise components. Moreover, we provide an intuitive illustration to understand this phenomenon using the concept of landscape representation. Mathematically, we analytically derive a trade-off relation that agrees well with simulations. Our results uncover a new property of noise in signaling processes, deepening our understanding of noise control and proposing a new perspective in designing signaling network.

## Introduction

Signaling processes are important for physical and biological networks. For example, in living systems, vital signaling processes include the transmission of environmental nutrient concentrations in bacteria chemotaxis [1], and the transmission of positional information in the development of multicellular organisms (morphogenesis) [2–5]. However, fluctuations inevitably affect the performance of signal transduction [6,7]. Over the years, mechanisms have been proposed to suppress the stochastic fluctuation in signaling networks, such as feedback control [8–11], spatiotemporal averaging [12,13], or integration of multiple signals [8,14–16].

To understand the mechanism of noise control and the properties of fluctuations, landscape representation provides an intuitive picture, in which the steady states of a system are mapped to local minima of a potential well in the state space [17–19]. And the width of the potential well is, to the first order, positively correlated with the relaxation timescale of the respective state steady.

However, it remains elusive whether the width of a potential well, i.e., the relaxation timescale, is positively or negatively correlated with noise in biological signaling systems. On the one hand, it has been proposed that long-timescale systems can average out fluctuations [10,12,13], resulting in a negative correlation between the width of the potential well and noise. On the other hand, long-timescale systems are proposed to suffer an accumulation of fluctuations [20,21], leading to a positive correlation, which is in consistent with the picture in thermodynamics.

A possible explanation to this puzzling question lies on the possibility that fluctuations from different sources could affect the noise differently in signaling processes [7,22–29]. While decomposing noise into extrinsic and intrinsic components is widely applied, its definition is slightly different depending on context [7,22,23]. For signaling systems, a natural decomposition is to dissect the noise into an “upstream component” that is originated from the fluctuation of the upstream input signal, and a “downstream component” that emerges from the stochasticity of the signaling process.

Here, we investigate the noise of signaling systems resulting from the upstream signal fluctuation and the downstream stochastic fluctuation, and discover that the control of the two noise components is opposite in terms of the width of the potential well. A wider potential well, corresponding to a longer timescale, can buffer upstream fluctuation, while accumulate downstream fluctuation. This contrariety leads to a fundamental trade-off in controlling the upstream and downstream noise. We verify that this trade-off relation also exists in real biological signaling networks, including a fold-change detection circuit [30] and the p53 activation signaling system [9]. Moreover, we provide an analytical trade-off relation at the ensemble level, which aligns well with numerical simulations. Interestingly, we find that a large timescale separation is detrimental for noise suppression in signaling systems.

## Results

### Modelling signaling systems with upstream and downstream fluctuation

To investigate how different noise components are affected by timescales, we first focus on a simplified signaling network. As illustrated in Fig. 1(a), we model a *N*-node network, which responds to an upstream signal *F*. Usually, the dynamics of such a system can be described by a set of nonlinear ordinary differential equations:

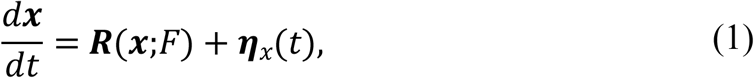

where ***x*** = (*x*_1_,…,*x*_*n*_)^*T*^ represents the state vector of the network, ***R*** = (*R*_1_,…*R*_*n*_)^*T*^ is the nonlinear regulatory function, *F* = *F*_0_ + *η*_*F*_(*t*) is the noisy upstream signal with a fluctuatio *η*_*F*_, *F*_0_= ⟨*F*⟩ is the average signal, ⟨…⟩ denotes temporal average, and ***η***_*x*_ = (*η*_*x*1,_…,*η*_*xn*_)^*T*^ is the downstream fluctuation originating from the stochastic reaction of the network. As a first step toward understanding the noise properties following the spirit of perturbation theory [28,31], we assume that the signaling system operates at a steady state and consider its dynamics near a stable fixed point ***x***_0_ [28]. Approximately, Eq. (7) becomes:

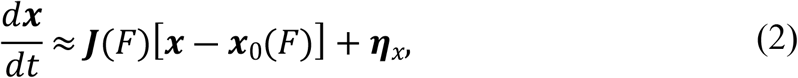

where ***J*** is the Jacobian at the stable fixed point ***x***_0_, i.e., 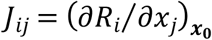. Assuming the fluctuation of the signal (*η*_*F*_) is small, Eq. (2) can be further approximated as:

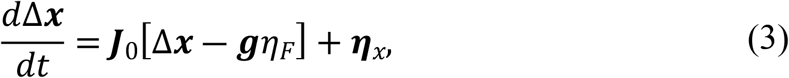

where Δ***x*** = ***x*** ― ***x***_0_(*F*_0_), ***J***_0_ = ***J***(*F*_0_), and ***g*** = *d****x***_0_(*F*_0_) *dF*_0_ represents the sensitivity (gain) [Fig. 1(b)].

**FIG. 1.**
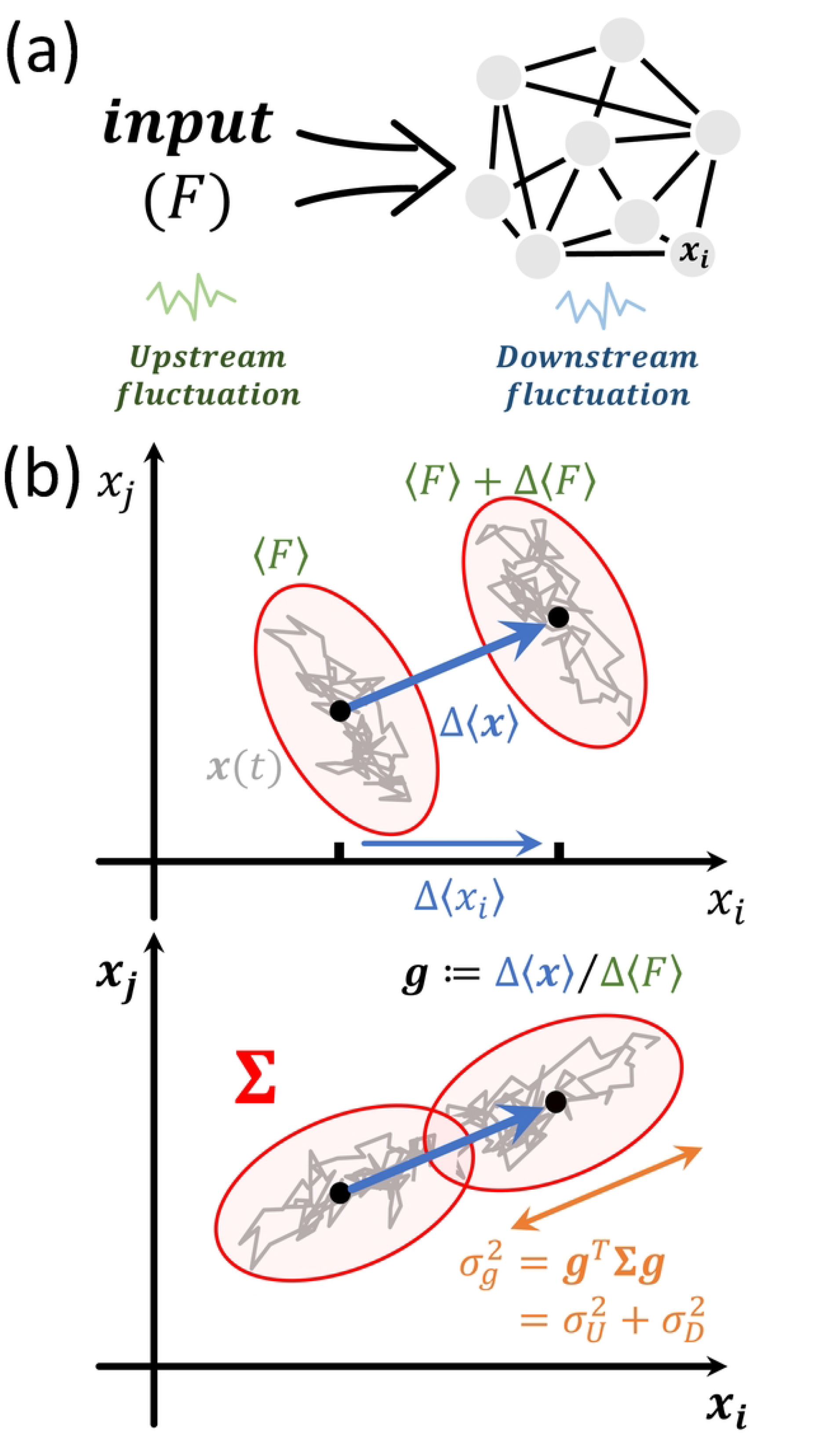
Schematic of the stochastic response of a signaling system. (a) In our model, a signaling system is modeled as a stochastic network receiving an upstream signal *F*, which *per se* is noisy. The activity of the node *i* is denoted as *x*_*i*_. (b) In the phase space, the effective region spread by the stochastic trajectory (gray line) of the system under a certain signal *F* is determined by the covariance matrix **Σ**, illustrated by the red ellipse. This region has a shift Δ⟨***x***⟩ (blue arrow) when the average input signal is varied by Δ ⟨*F*⟩, defining a sensitivity vector (or gain) ***g***. The noise along the ***g*** direction, denoted as 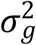, is vital for signal transduction, since a large 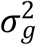 (bottom) could lead to an overlap of trajectories under different signals comparing to a small 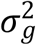 (top). The noise along the ***g*** direction is given as 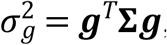, which has two components 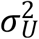 and 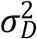 contributed by the upstream fluctuation denoting the fluctuation of the input signal and the downstream fluctuation representing the stochasticity of the network, respectively.

Throughout the analysis, we assume the upstream fluctuation *η*_*F*_ has a correlation time of *τ*_*F*_, i.e., 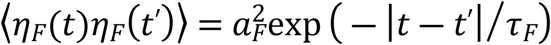, where *a*_*F*_ is the fluctuation amplitude, and the downstream fluctuation ***η***_*x*_ is independent white noise, i.e., 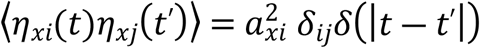, where *a*_*xi*_ is the fluctuation amplitude along the *x*_*i*_ direction and assumed to be constant. Although generally the downstream fluctuation could depend on the instantaneous state of the system or even other dynamic parameters of the system (multiplicative noise), there is no general relation between *a*_*xi*_ and ***x***_**0**_ [17]. For simplification, we assume *a*_*xi*_ is constant, which is equivalent to probe how susceptible the system is to the downstream fluctuation. Hence, all terms in the right-hand side of Eq. (3) are to the same order.

The noise of this dynamical system can be described by the covariance matrix at the steady state **Σ = ⟨Δ*x*Δ*x***^*T*^⟩. In addition to the noise in *x*_*i*_ direction 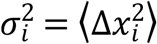, we also consider the noise along the ***g*** direction denoted as 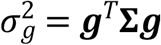, which quantifies how distinguishable two different input signals are form given stochastic trajectories. For example, in Fig. 1(b), a small difference of the input level is relatively easy to be distinguished in the system represented by the top panel but not the bottom panel.

### An intuitive trade-off between upstream and downstream noise in 1-dimensional systems

To gain an intuitive picture on the difference between upstream and downstream fluctuations, we first study a one-dimensional system in which Eq. (3) becomes:

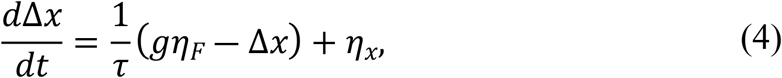

where *τ* is the timescale of the system. Since Eq. (4) can be described as a gradient system: *d*Δ*x dt* = ― ∂*U* ∂Δ*x* + *η*_*x*_ where 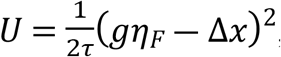, the timescale *τ* also represents the width of the potential well.

This linear system can be analytically solved as:

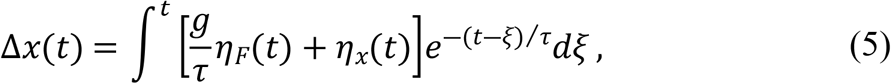

and the noise follows:

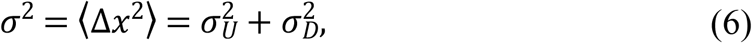

where 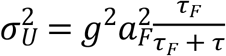 and 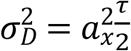 are the two noise components originated from the upstream fluctuation and downstream fluctuation, respectively. Equivalently, the susceptibility to the upstream and downstream fluctuation are given by 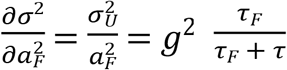 and 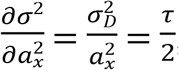, respectively.

This result indicates that the dependence of the susceptibility to upstream fluctuation and downstream fluctuation on the width of potential well *τ* is opposite to each other: 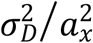 is negatively correlated with *τ*, while 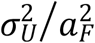 is positively correlated with *τ*. In other words, there is a trade-off in controlling the two noise components.

Intuitively, upstream fluctuations and downstream fluctuations could be mapped to the fluctuation of the potential well *per se* (the “bowl” in Fig. 2) and the state variable (the “ball” in Fig. 2), respectively. A shorter relaxation timescale corresponds to a narrower, therefore sharper, potential well, which means that the state of the system pursues closely to the valley. As a result, the fluctuation of the “bowl” will completely propagate to the “ball”, resulting in high susceptibility to the upstream fluctuation. On the other hand, the downstream fluctuation of the system is strictly confined since any small fluctuation of the “ball” relative to the “bowl” will be recovered in a short timescale, being unlikely for accumulation. If the downstream fluctuation also has a nonzero correlation time *τ*_*x*_, the downstream noise component becomes: 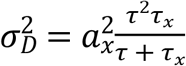, which is still positively correlated with the width of the potential well *τ*.

**FIG 2.**
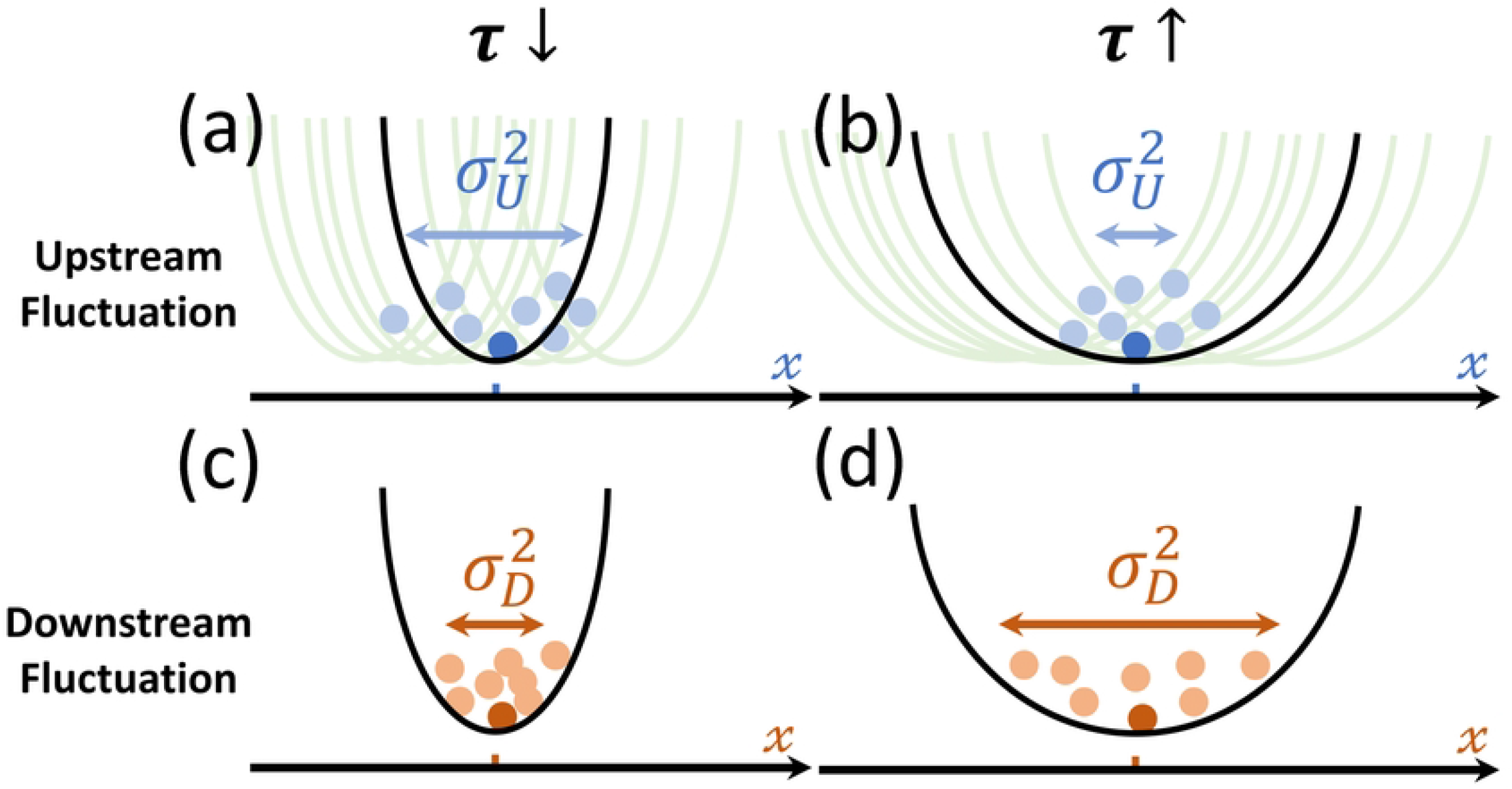
Illustration of the trade-off between the upstream noise and the downstream noise on the potential well. The width of a potential well is positively correlated with the timescale *τ* of the dynamical system. (a, b). On the one hand, signaling systems are more sensitive to the upstream fluctuation (the fluctuation of the well, denoted as light green curve) with a narrower potential well (a) comparing to a wider one (b). For example, a “ball” in a wider, therefore flatter, “bowl” is less susceptible to the fluctuation of the “bowl” itself (under the overdamped limit). (c, d) On the other hand, signaling systems are less susceptible to downstream fluctuation (such as thermal fluctuation) with a narrower potential well (c) comparing to a wider one (d), since a narrow well means a random displacement can be recovered rapidly and less likely to accumulate.

### A general trade-off relation in high-dimensional and nonlinear systems

While one-dimensional systems generally follow gradient dynamics, high-dimensional systems may not. To investigate the generality of this trade-off, especially in non-gradient systems, we numerically simulate the dynamics [Eq. (3)] of high dimensional systems. Specifically, we choose a set of parameters for a two-node network and a ten-node network, respectively [Figs. 3 (a) and (b)], and evaluate the noise along the ***g*** direction 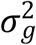 and the noise of a single node 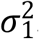. When tuning the overall timescale of the system via setting ***J***_**0**_***→J***_**0**_/*γ*, an opposite trend for the upstream noise and downstream noise appears for both networks [Figs. 3 (c-f)], indicating the trade-off relation also exists in high dimensional systems. As a result, the total noise exhibits a nonmonotonic dependence on the overall timescale.

**Fig.3.**
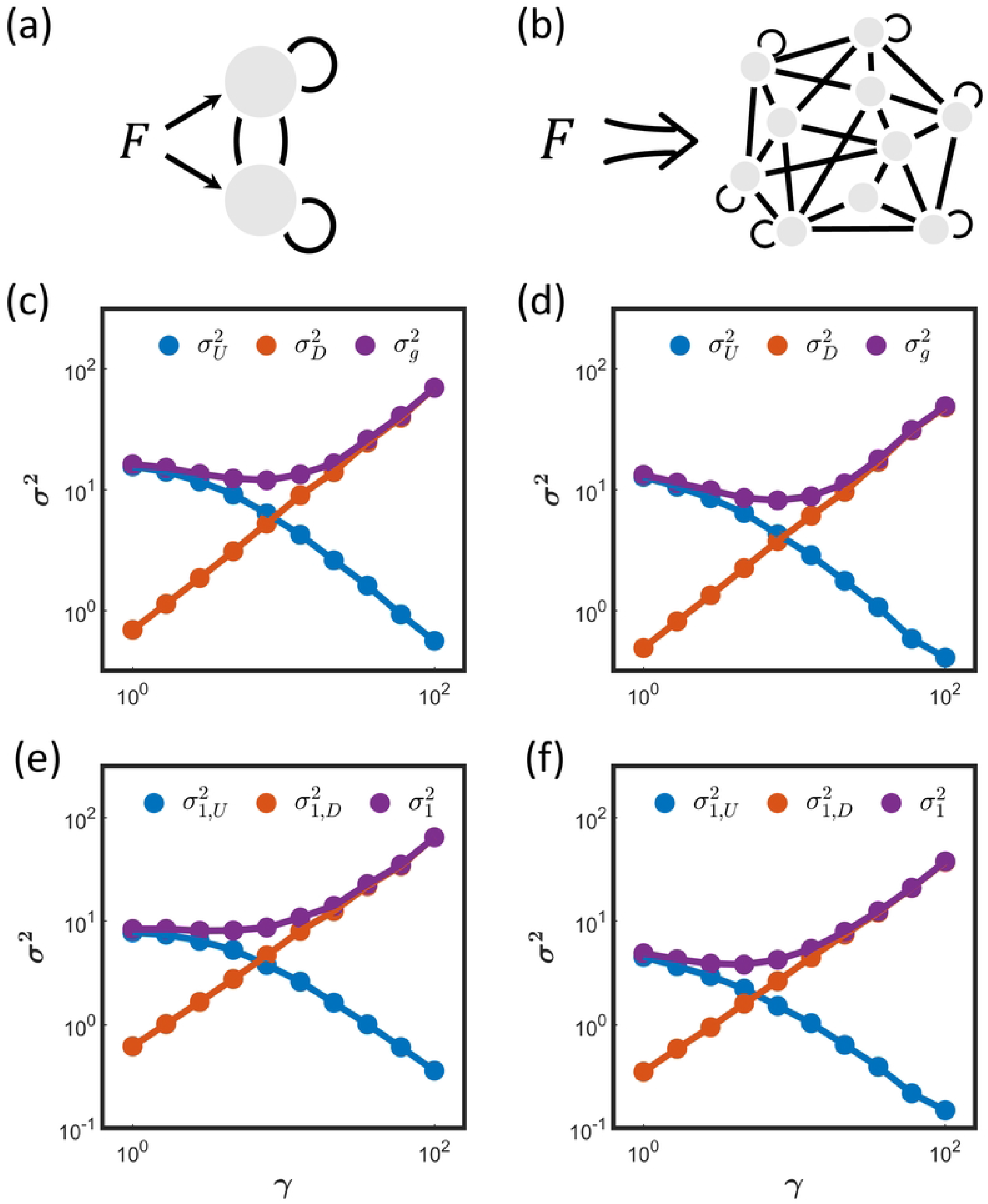
The trade-off between upstream and downstream noise in high dimensional systems. (a, b) A two-node network (a) and a ten-node network (b) are used as an illustration. (c, d) In both networks, when tuning the overall timescale (*γ*), the noise from upstream fluctuation 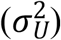 and downstream fluctuation 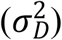 exhibit opposite trends, indicating a potential trade-off. The total noise 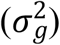 follow the linear decomposition and shows a nonmonotonic trend. (e, f) The noise of an arbitrary node 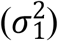 for both networks follows the same trade-off relation.

To test whether the derived trade-off relation exists in real biological signaling networks with nonlinearity, we numerically simulate a typy-1 incoherent feedforward loop circuit capable for fold-change detection [30] and the UV-induced p53 activation pathway [9]. Apart from the nonlinearity, we use multiplicative downstream fluctuations for both models following the literature [30] (see S1 Appendix for details).

In biological systems, usually only a subset of node in the network is used to transmit signal. Therefore, in these models, we only consider the noise of the output node. Our results show that, in both models, the response noise can be linearly decomposed into the upstream component and downstream component following the form 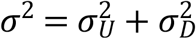 [Fig. 4]. Moreover, the dependence of the two components on the overall timescale of the system is inversed, similar to the theorical result. These results suggest the generality of the proposed trade-off relation in real biological systems.

**Figure 4.**
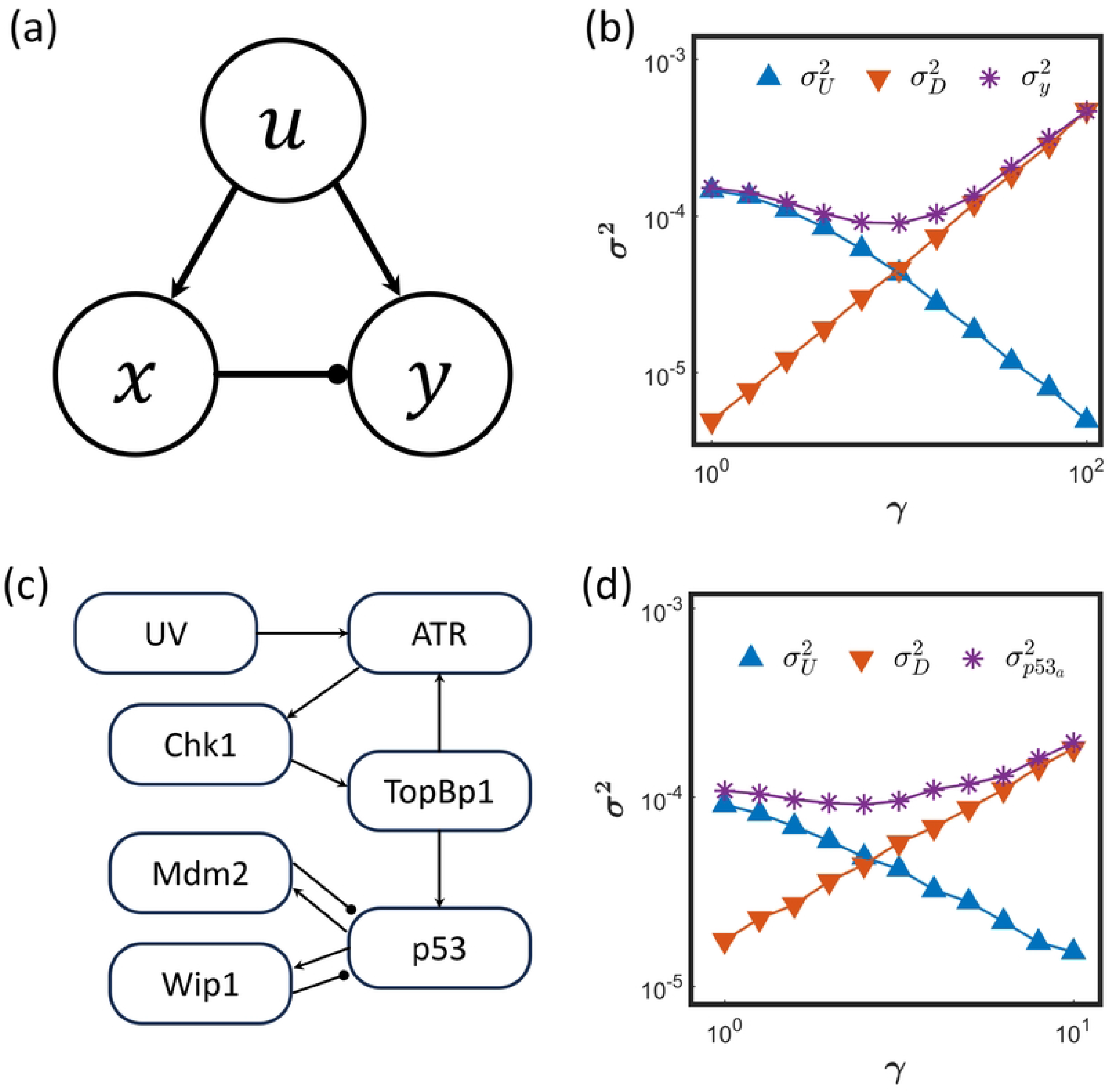
The trade-off between upstream and downstream noise component is verified in two representative nonlinear biological models: fold-change detection circuit (a, b) and p53 signaling systems (c, d). (a) A type 1 incoherent feedforward loop for fold change detection. (b) The steady state noise of the response node *y* can be decomposed into two components originated from upstream and downstream fluctuation, respectively. A trade-off between the two noise components is observed when tuning the timescale of the system. *γ* denotes the fold change of the overall timescale of the system. (c) The p53 activation signaling pathway containing 7 nodes. (d) The steady state noise of the response, active p53, can also be decomposed into two components originated from upstream and downstream fluctuation, respectively. The trade-off is also observed.

### Ensemble level trade-off and the lower bound

So far, our results suggest a trade-off between upstream and downstream noise in tuning the timescale at the single network level. An immediate question is whether this trade-off relation appears at the ensemble level. To test it, we randomly choose 20,000 different parameter sets (***J***_0_ and ***g***) for 2-node networks and 10-node networks, respectively (see S1 Appendix for details). And *a*_*xi*_ = *a*_*x*_ and *a*_*F*_ are fixed throughout the simulation. We choose a time unit such that *τ*_*F*_ = 1, and ***g*** is normalized such that |***g***| = 1. Under these constraints, the results from different parameter sets are comparable. Notably, the noise of a specific node 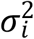 depends on its sensitivity *g*_*i*_ to the upstream signal, which could vary between different parameter sets, hence it is not comparable at the ensemble level. Therefore, we focus on the ***g***-direction noise.

In the simulation results, we observe a global trade-off between the upstream noise 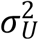 and the downstream noise 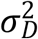 (Fig. 5). All the data points appear above a clear boundary (Fig. 5). And consistent with the physics intuition (Fig. 2), at the ensemble level, an increase of the “effective width of potential well” *τ*, defined as the negative inverse of the eigenvalue of the Jacobian matrix ***J***_**0**_ (see S1 Appendix for details), can attenuate the noise from upstream fluctuation, and amplify the noise from downstream fluctuation, resulting in a decrease of 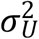 and increase of 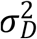. The examples used in Fig. 3 is shown as the yellow curve in the figure, indicating the “single network trade-off relation”. This result also indicates that the minimal value of total noise can be achieved when the upstream and downstream noise balance in an optimal timescale [the dark red arrow in Fig. 5 (a) and (b)].

**FIG. 5:**
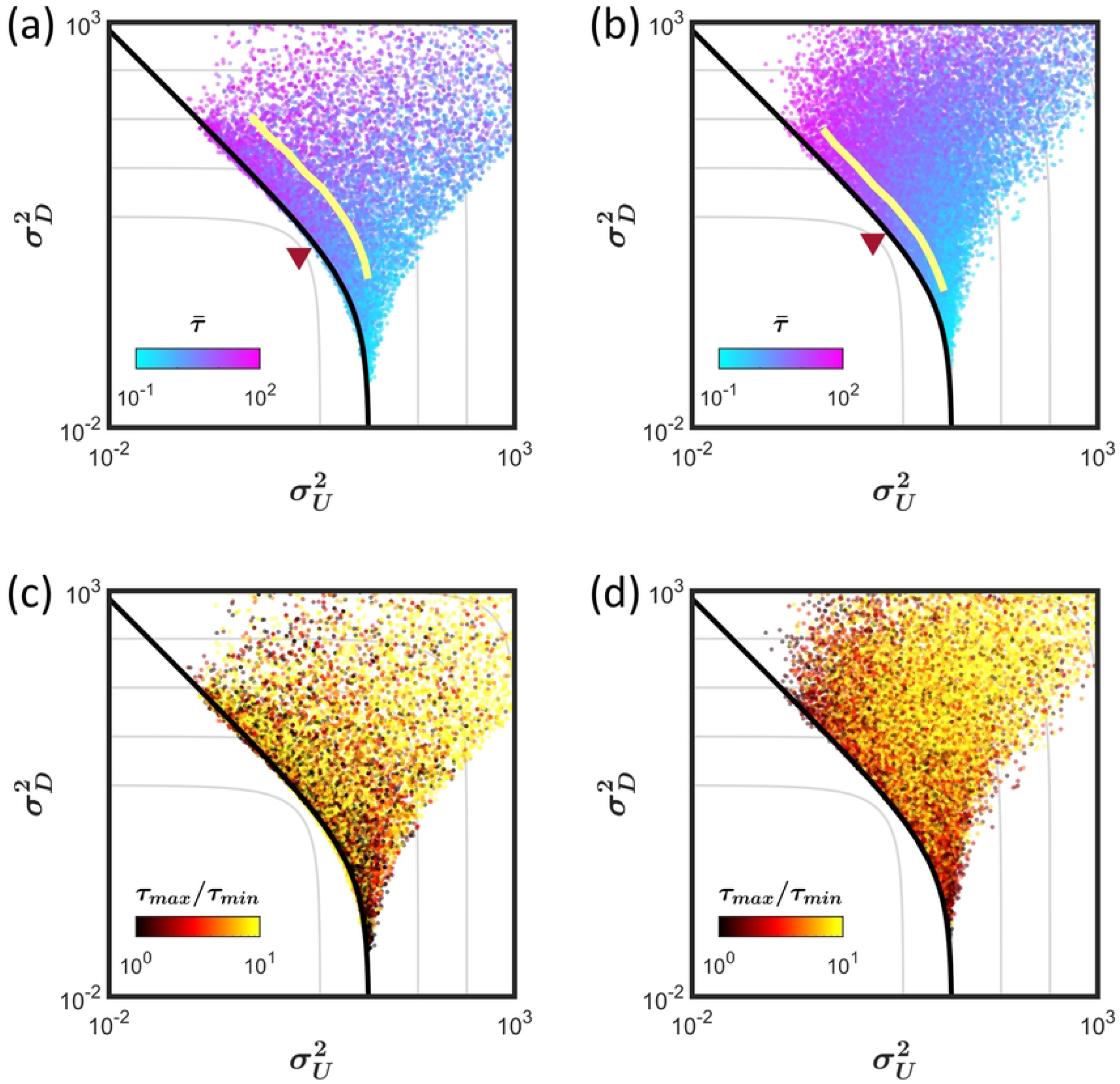
The trade-off between noise from upstream fluctuation and downstream fluctuation at the ensemble level. (a, b) By sampling the parameters of the two-node network (a) and ten-node network (b), the simulation results lie above a clear curve on the bottom-left (black curves), which are analytically derived. The dot color denotes the geometric mean of all timescales of the system 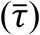 in each specific parameter set. Gray curves represent the contour of the total noise 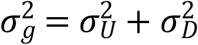. To achieve the minimal total noise, the timescale of the system should reach an optimal value (red triangle). The individual level trade-off in Figs. 3 (c) and (d) are shown as yellow curves. (c, d) The dot color denotes the extent of timescale separation (*τ*_*max*_ *τ*_*min*_), which increases the overall noise for both components.

Mathematically, finding the lower bound of the trade-off relation is equivalent to minimizing 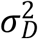 for given 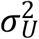 for the dynamic system [Eq. (3)]. We find that all timescales being equal is a condition for minimizing the Lagrangian function 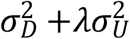 (see S1 Appendix for details). Following this condition, we find a general, dimension independent, trade-off relation for the two noise components:

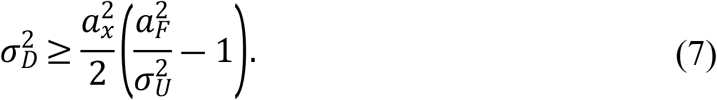

The curve showing Eq. 7 aligns well with the bottom-left boundary of the data points for all the simulation systems we tested (Fig. 5), supporting the dimension independence.

Surprisingly, if the system shows a timescale separation, quantified by a large ratio between the maximal timescale and the minimal timescale, i.e., *τ*_*max*_ *τ*_*min*_(see S1 Appendix for details), the resulting 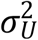 and 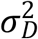 will shift away from the lower bound [Fig. 5 (c) and (d)], suggesting that the timescale separation could be detrimental for noise control in signaling. A possibility is that when the system shows timescale separation, it is susceptible to the upstream fluctuation in the fast directions and it accumulates the downstream fluctuation in the slow directions. Moreover, any small perturbation away from the steady state will project onto the slow manifold in a short timescale along the fast dimension, which could amplify the absolute noise amplitude if this projection is not orthogonal (Fig. S1).

## Discussion

To providing a clear intuition, we use linear approximation and additive noise in deriving the ensemble trade-off relation [Eq. (7), Fig. 5]. Although the trade-off exists for nonlinear model with multiplicative noise in single network level (Fig. 4), deriving the lower bound of the ensemble level trade-off under this general condition is challenging and require further studies with advanced technique. Nonetheless, the image provided here is clear: in a small noise condition, linear approximation is sufficient, and the susceptibility to upstream and downstream fluctuation follows the trade-off relation we derived.

Notably, the downstream fluctuation discussed here is different from the intrinsic noise defined in some literatures, which usually refers in specific to Poisson noise [24,32]. Indeed, there is no guarantee that the downstream fluctuation of a complex signaling systems follows a Poisson process. However, under some certain extreme conditions, if the downstream fluctuation is dominated by the Poisson noise, i.e., 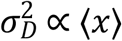, and therefore it could be independent to the timescale, the width of the potential well will *negatively* correlate with the noise 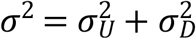 because of the negative correlation between 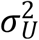 and *τ*, i.e., a wider potential well will, counterintuitively, results in lower noise, which is still against the well-known image of thermal-fluctuation in physics.

While in some biological systems, a well-defined component is regard as the output node (Fig. 4), the ***g***-direction is the most informative dimension in information theory (Fig. 1). We confirm that the single network level trade-off is satisfied for both the ***g***-direction noise and natural direction noise (Fig. 3). However, for the ensemble comparison, only the ***g*-**direction noise is comparable between parameter sets. Moreover, the ***g***-direction noise is closely related to the information transmission capacity of the signaling pathway.

For general signaling systems, the dimension of the parameter space is usually high, our result suggests that only two dimensions, the overall timescale and the extent of timescale separation, are stiff for the noise. In short, tuning the overall timescale adjusts the susceptibility to upstream fluctuation and downstream fluctuation which is, however, constraint by a trade-off; tuning the extent of timescale separation can control the two noise components toward the same direction. This is in consistent with the concept that biological systems are usually sloppy [33,34].

Finally, noise control in signaling processes is vital for many biological functions. The discovered general trade-off in controlling the noise from the upstream and downstream fluctuation could be helpful in the rational design of signaling networks for synthetic biology.

## Supporting information

**S1 Appendix. Supporting information.** In this note, we provide the details of parameter sampling in numerical simulations (Figs. 3 and 5), and the details of the biological model regarding Fig. 4. Also, we provide a detail mathematical derivation of the lower bound Eq. (7) in Fig. 5. And we give an explanation of the effect of timescale separation on noise.

## Acknowledgements

This project is supported by the National Natural Science Foundation of China 32271293 and 11875076. The numerical simulation was performed on the High Performance Computing Platform of the Center for Life Sciences, Peking University.

